# Integrative modeling in the age of machine learning: a summary of HADDOCK strategies in CAPRI rounds 47-55

**DOI:** 10.1101/2024.09.16.613212

**Authors:** Victor Reys, Marco Giulini, Vlad Cojocaru, Anna Engel, Xiaotong Xu, Jorge Roel-Touris, Cunliang Geng, Francesco Ambrosetti, Brian Jiménez-García, Zuzana Jandova, Panagiotis I. Koukos, Charlotte van Noort, Joao M. C. Teixeira, Siri C. van Keulen, Manon Réau, Rodrigo V. Honorato, Alexandre M.J.J. Bonvin

## Abstract

The HADDOCK team participated in CAPRI rounds 47-55 as both server, manual predictor, and scorers. Throughout these CAPRI rounds, we used a plethora of computational strategies to predict the structure of protein complexes. Of the 10 targets comprising 24 interfaces, we achieved acceptable or better models for 3 targets in the human category and 1 in the server category. Our performance in the scoring challenge was slightly better, with our simple scoring protocol being the only one capable of identifying an acceptable model for Target 234. This result highlights the robustness of the simple, fully physics-based HADDOCK scoring function, especially when applied to highly flexible antibody-antigen complexes. Inspired by the significant advances in machine learning for structural biology and the dramatic improvement in our success rates after the public release of Alphafold2, we identify the integration of classical approaches like HADDOCK with AI-driven structure prediction methods as a key strategy for improving the accuracy of model generation and scoring.

## 1 Introduction

The last few years have witnessed a profound revolution in computational structural biology, with deep learning models exploiting the huge amount of experimentally resolved structures available in the Protein Data Bank (PDB) [1]. Together with the Critical Assessment of protein Structure Prediction (CASP), the Critical Assessment of PRediction of Interactions (CAPRI) experiment has played a pivotal role in evaluating the methodological progress and identifying the remaining challenges within the field of biomolecular complex structure prediction. CAPRI rounds 47-55 represent a time travel through the deep learning revolution, with the first target (Target 160) dating back to early 2019 and the last one (Target 234) being released in late 2023.

We participated in CAPRI rounds 47-55 (excluding the CASP-CAPRI rounds 50 [2] and 54 [3] and round 52), contributing to all three categories—server, manual/human, and scoring—using HADDOCK [4], our data-driven integrative modeling platform, under various team names (HADDOCK for the server, and Bonvin and Giulini as human teams). Specifically, we used HADDOCK versions 2.4 [5] and 3.0 (beta5), converting the information we could find about the complexes to be predicted into ambiguous restraints to guide the modeling process. It is however important to note that, while good information allows to properly drive the docking towards native solutions, incorrect or incomplete input information might cause the docking to fail, as a wrong region of the conformational space will be sampled.

In the Methods section, we describe the strategy followed and information used by target, separating them into two groups based on their release date, namely before or after the availability of the public version of AlphaFold2 [6, 7]. We then discuss our performance according to the CAPRI evaluation results.

## 2 Methods

Here we briefly describe the procedures we followed to model the targets in CAPRI rounds 47-55. The diverse nature of the targets and the different release times make it impossible to define a unique procedure for all of them, but we can highlight the major steps common to all the modeling efforts. Subsequently, the details of the strategy followed for each target are presented more in depth.

### 2.1 General Procedure

The first step of the modeling procedure consists of retrieving or generating structural models of the unbound partners, if not already provided. For CAPRI rounds 47-51 template-based homology modeling strategies were followed [4, 8, 9]. Instead, for CAPRI rounds 53 and 55 AlphaFold2 [6] and another deep learning-based structure predictor [10] were used for this task.

After modeling the input structures, interface information was obtained using biological knowledge or literature data. As an example of the former, complementarity-determining region (CDR) residues of antibodies are known to be the regions involved in antigen recognition. Literature data include but are not limited to interfaces between homologous complexes, mutagenesis or other experimental data, and bioinformatic predictions. Target 160 (discussed below) represents an unusual exception.

Once provided with input structures and interface information translated into ambiguous restraints, HADDOCK was used to dock the molecules together following its standard pipeline, which consists of rigid-body docking followed by a semi-flexible refinement by molecular dynamics in torsion angle space, finalized by an energy minimization step. The HADDOCK scoring function proper to each stage [11, 12] is used to score the generated models.

Given the challenging nature of the targets (see Table 1), we aimed at having a diverse set of models in our submission ensembles. This diversity was typically achieved by performing a Fraction of Common Contacts (FCC) clustering [13] step after the refinement and usually selecting a representative from the top 5 ranked clusters.

**Table 1.** Prediction and scoring results obtained by the HADDOCK server team and by the Bonvin and Giulini manual teams for all targets of the five CAPRI rounds. All the evaluated interfaces are shown. We report the number of high(***), medium (**) and acceptable (*) quality models based on CAPRI criteria. The best DockQ value obtained for each set of models is also reported between square brackets. All the targets are labelled as “difficult” to predict with the exception of Target 232 (labelled as “easy”).

| Target | Target Category | Strategy | Interface | Server<br>*** / ** / * | Human<br>*** / ** / * | Scoring<br>***/**/* |
| --- | --- | --- | --- | --- | --- | --- |
| 160 | Nanobody - Protein | picture - driven | T160.1 | 0 / 0 / 0 [0.02] | 0 / 0 / 0 [0.03] | 0 / 0 / 0 [0.02] |
|  |  |  | T160.2 | 0 / 0 / 0 [0.04] | 0 / 0 / 0 [0.03] | 0 / 0 / 0 [0.05] |
|  |  |  | T160.3 | 0 / 0 / 0 [0.19] | 0 / 0 / 0 [0.19] | 0 / 0 / 0 [0.19] |
|  |  |  | T160.4 | 0 / 0 / 0 [0.01] | 0 / 0 / 0 [0.01] | 0 / 0 / 0 [0.01] |
|  |  |  | T160.5 | 0 / 0 / 0 [0.03] | 0 / 0 / 0 [0.05] | 0 / 0 / 0 [0.03] |
|  |  |  | T160.6 | 0 / 0 / 0 [0.10] | 0 / 0 / 0 [0.10] | 0 / 0 / 0 [0.10] |
|  |  |  | T160.7 | 0 / 0 / 0 [0.14] | 0 / 0 / 0 [0.14] | 0 / 0 / 0 [0.14] |
|  |  |  | T160.8 | 0 / 0 / 0 [0.02] | 0 / 0 / 0 [0.02] | 0 / 0 / 0 [0.02] |
|  |  |  | T160.9 | 0 / 0 / 0 [0.13] | 0 / 0 / 0 [0.13] | 0 / 0 / 0 [0.13] |
| 161 | Protein - Protein | Template - based | T161.1 | 0 / 0 / 0 [0.02] | 0 / 0 / 0 [0.04] | 0 / 0 / 0 [0.02] |
| 162 | Protein - Protein | Template - based | T162.1 | 0 / 0 / 0 [0.04] | 0 / 0 / 0 [0.03] | 0 / 0 / 0 [0.03] |
| 163 | Protein - Protein | Information-driven | T163.1 | 0 / 0 / 0 [0.02] | 0 / 0 / 0 [0.02] | 0 / 0 / 0 [0.02] |
|  |  |  | T163.2 | 0 / 0 / 0 [0.02] | 0 / 0 / 0 [0.04] | 0 / 0 / 0 [0.03] |
|  |  |  | T163.3 | 0 / 0 / 0 [0.01] | 0 / 0 / 0 [0.01] | 0 / 0 / 0 [0.01] |
|  |  |  | T163.4 | 0 / 0 / 0 [0.00] | 0 / 0 / 0 [0.00] | 0 / 0 / 0 [0.00] |
|  |  |  | T163.5 | 0 / 0 / 0 [0.00] | 0 / 0 / 0 [0.00] | 0 / 0 / 0 [0.00] |
| 187 | Protein-Protein | ML - driven | T187.1 | 0 / 0 / 0 | 0 / 0 / 0 [0.02] | 0 / 0 / 0 [0.01] |
|  |  |  | T187.2 | 0 / 0 / 0 | 0 / 0 / 4 [0.40] | 0 / 0 / 1 [0.45] |
| 188 | Protein-Protein-DNA | Information-driven | T188.1 | 0 / 0 / 0 [0.01] | 0 / 0 / 0 [0.06] | 0 / 0 / 0 [0.06] |
|  |  |  | T188.2 | 0 / 0 / 0 [0.28] | 0 / 0 / 0 [0.28] | 0 / 0 / 0 [0.27] |
| 231 | Antibody - peptide | Information-driven | T231.1 | 0 / 0 / 0 [0.53] | 0 / 0 / 2 [0.64] | 0 / 1 / 3 [0.77] |
| 232 | Protein - protein | ML - driven | T232.1 | 0 / 4 / 0 [0.78] | 0 / 2 / 2 [0.76] | 0 / 10 / 0 [0.80] |
| 233 | Antibody - MHC | information-driven | T233.1 | 0 / 0 / 0 [0.07] | 0 / 0 / 0 [0.08] | 0 / 0 / 0 [0.05] |
| 234 | Antibody - MHC | information-driven | T234.1 | 0 / 0 / 0 [0.06] | 0 / 0 / 0 [0.22] | 0 / 0 / 1 [0.27] |

For the scoring experiment, we followed the standard HADDOCK scoring recipe [11, 12], in which all missing atoms are first built followed by a short energy minimization (50 steps of steepest descent). No restraints representing external information are used at this scoring stage. The models are then clustered based on the fraction of common contacts using a 0.6 cutoff and the resulting clusters ranked based on the average score of the top 4 members. The top ranking model of the top 10 clusters are selected for submission, after visual inspection, making sure there are no knots or intertwined regions at the interface(s) and if artefacts are found, instead the next model of the cluster is selected. An example HADDOCK 3.0 scoring protocol is provided in the Supplementary Information.

### 2.2 Pre-AlphaFold2 Targets

#### 2.2.1 Target 160

This target consists of the assembly of 6 domains from the S-layer protein SAP of *Bacillus anthracis* interacting with two nanobodies. Due to an internet leakage, this round is also known as “The Twitter target” in the BonvinLab history. Indeed, while this complex was presented at a meeting, a picture showing the complex in the background was posted on social media. In the search for suitable information to drive the docking, this picture was used to derive interaction restraints by measuring on a paper-printed figure distances and angles between the center of mass of the 6 SAP domains to be modeled (Fig 1.A) and converting those to angstrom using a scaling factor based on the estimated size of the domains. As those domains form in a planar arrangement, additional restraints to a plane consisting of artificial beads were defined to ensure planarity of the sampled solutions. Those were defined as ambiguous restraints between selected CA atoms and all the beads defining the plane. As backbone atoms of the individual domains were provided, the side-chains were modeled with SCWRL [8]. Nanobodies were homology-modeled based on templates 3AB0 [14] and 5MJE [15], for nb684 and nb694 respectively, using KotaiAB [16]. Additional conformational sampling of the CDR loops was performed with Modeller [17]. Residues of CDR loops were predicted and filtered for their solvent accessibility and later used to drive the docking onto the pre-established complex of SAP. An additional repulsion restraint of 50 Åwas added between the two nanobody centers of mass, making sure to sample solutions where the two nanobodies are not interacting.

**Figure 1.**
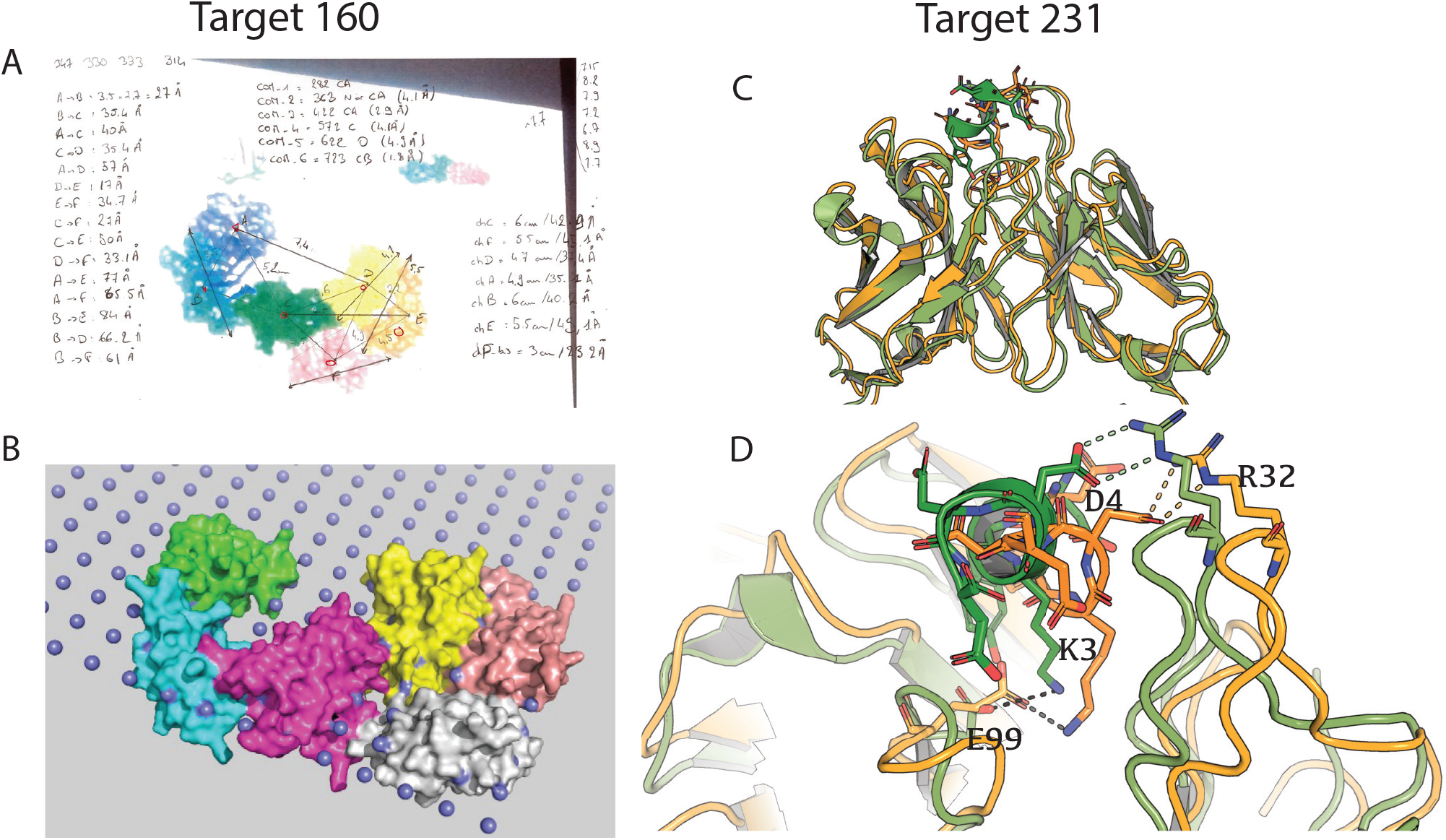
A) Visual representation of the picture-driven docking procedure. The picture posted on twitter was projected on a white board and distances between center of mass were drawn by hand and later converted into angstrom after estimating a scaling factor by measuring various domain sizes in PyMol [34], allowing to obtain restraints to guide the docking. B) Restraints to a plane were added to ensure planarity of docking solutions as this complex was known to 2D-self assemble and sit flat on the bacterial surface layer. This information was used to guide the 6-body docking of the SAP domains, which while not being correct visually reproduces the Twitter image. C) Our top ranked prediction model for target 231 (orange) superimposed onto the crystal structure of the reference complex (green) (PDB ID 8RMO) [33]. D) Despite the lack of helicity in the peptide conformation, major interactions are correctly identified, namely hydrogen bondings between Aspartic acid 4 (D4) from peptide to Arginine 32 (R32) of the light chain and the salt-bridge between Lysine 3 (K3) from peptide with Glutamic acid (E99) of the heavy chain.

#### 2.2.2 Targets 161 and 162

The system for both targets involves the antitoxin RnlA from *E. coli*, as a homodimer for target 161 and bound to the toxin RnlB for target 162. For target 161, the homodimeric form of RnlA must be predicted. To this aim, available structural homologues were extracted from the PDB, where 4I8O [18] was used to build a structural model of the DBD domain with Modeller [17] and 5HY3 [19] to generate a model of the N-repeated domain. Interface restraints derived from the template 5HY3 were used to guide the 4-body docking (the two domains of the monomer were treated as separate entities to allow for more conformational changes). For target 162, the backbone of RnlB was provided, therefore HADDOCK was used to build the atomic coordinates for the missing side-chains residues. For RnlA, we used our top1 target 161 model to extract connectivity restraints. Each structural domain from RnlA was separated into a single chain, removing flexible linkers between them, and residues in contact with the toxin in 5HY3 defined as active, defining the binding site for RnlB. For the RnlB domains, no information could be retrieved, therefore solvent-accessible surface residues were treated as passive in the definition of the restraints (e.g., they can but do not have to be at the interface).

#### 2.2.3 Target 163

Both SYCE2 and TEX12 models were obtained through homology modeling. SYCE2 was built using multiple tools [9,17,20] and refined with HADDOCK2.4, while TEX12 was modeled from two different PDB structures (chain A of 6HK8 and chain A of 6HK9) to capture conformational diversity. The RMSD between the two conformations is 1.5 Å.

Restraints were applied based on the interaction pattern proposed by Davies *et al*. (2012) [21], aiming to model the SYCE2-TEX12 hetero-complex that exists at the core of the synaptonemal complex. Although the generated models satisfied the restraints, no acceptable model was produced. Interestingly, the initial model by Davies *et al*. was later revised [22] to a completely different interaction pattern, where the N-terminal regions are interacting to form filaments, and the corresponding reference structure deposited with PDB ID 6R17.

### 2.3 Post-AlphaFold2 Targets

#### 2.3.1 Target 187

The TNPA protein was obtained using a local version of AlphaFold2.1. The DNA was built in a B-form using 3D-DART [23]. In the web server modeling, all solvent accessible arginine residues are restrained to be in contact with the C1/C9 DNA atoms through ambiguous restraints excluding the DNA ends (5 residues on each side).

In the human submission we used a slightly different setup, in which C2 symmetry and non-crystallographic symmetry are enforced between the two protein-DNA molecules. To allow for tighter solutions the intermolecular van der Waals energy was scaled down by a factor 0.001 and its weight set to 0.0 during the rigid-body docking phase. The desolvation energy weight was set to 0.0 for all stages of the HADDOCK2.4 workflow.

#### 2.3.2 Target 188

We used the same protein conformation as in Target 187. Here a five-body docking was performed, treating the N-terminal regions of the protein as independent domains with connectivity restraints [24] to ensure reasonable proximity of the backbone as the cutting point. In addition to symmetry restraints, several catalytic residues on the protein were coupled to Phosphate atoms of the DNA. Terminal Arginines NH groups were also restrained to be in contact with the Phosphate atoms of DNA. Last, N-terminal domains are coupled to the DNA.

#### 2.3.3 Target 231

We used PDB file 7BG1 [25] as starting point for the antibody structure, as the sequences are identical. For the FLAG peptide, we took the five conformations extracted from Alphafold2.3. Those models were subjected to a very short MD refinement (100 ps) in explicit water solvent using the OpenMM [26] module implemented in HADDOCK 3.0. An ensemble of 8 conformations was produced upon clustering the short MD trajectories. This strategy was not good, as the obtained peptide models do not show the 3_10_-helical conformation observed in the bound structure.

The visual inspection of the accessible surface around the antibody CDR region of PDB 7BG1 revealed the existence of a negatively charged pocket centered around Glutamic acid E99 from the antibody heavy chain. This residue was then paired to one of the two positively charged residues on the FLAG peptide (Lysine K3) with the aim of compensating the excess negative charge of the pocket. Lysine K8 was not considered here as it was the last amino acid of the FLAG peptide. The docking was performed with HADDOCK imposing this single contact restraint. The best 4 models from the top 25 clusters were then post-processed with a 10 ns MD simulation with OpenMM. Models were rescored based on their conformational stability with respect to the initial docking pose, by tracking the evolution of DockQ over time [27].

#### 2.3.4 Target 232

For this target we used AlphaFold2.3 and HADDOCK3 to model the complex. HADDOCK3 was first used together with restraints derived from the analysis of AlphaFold2.3 models, and also with information coming from the literature [28, 29]. Then, both server and human ensembles were created placing AF2 predictions, AF2-derived HADDOCK models and literature-derived HADDOCK models in the HADDOCK 2.4 refinement interface [5].

#### 2.3.5 Targets 233 and 234

Both targets 233 and 234 feature the interaction of a major histocompatibility complex (MHC) with an antibody. We used AlphaFold2.3 to model the MHC complexes (Human leukocyte antigen A with Beta 2 microglobulin), and we combined AlphaFold2.3 and Immunebuilder [10] predictions for the antibody modeling task, clustering the retrieved antibody structures based on the RMSD calculated on the H3 CDR loop after superimposition on the conserved region. As restraints, we defined part of the CDR residues on the antibody side as active in HADDOCK using proABC2 [30]. For the MHC molecule, we extracted the contacts found in the literature between these molecules and any antibody and complemented those with all the contacts found by AlphaFold2.3 on the multimeric predictions.

## 3 Results and Discussion

Table 1 shows the results obtained for the three CAPRI experiments. We obtained acceptable-or higher quality models for 3 and 1 of the 24 interfaces in the human and server predictions, which corresponds to success rates of 13 and 4%. As these numbers are heavily biased by the incorrect modeling for Target160, we can consider targets rather than interfaces, which brings our success rates to 30% and 10%, respectively. The performance is better in the scoring challenge, with a success rate of 17% (resp. 40%) considering interfaces (resp. different targets). If we consider only targets released in the post AlphaFold2 era, the success rates are much higher despite the objective difficulty of five of the six targets. Indeed, we obtain 17 %, 50 % and 67 % target-based success rates for server, human and scoring challenges, respectively.

AlphaFold’s unique capability to generate reliable structural models of protein complexes has fundamentally transformed the field of computational protein interaction prediction. When such prediction is wrong (for example in the case of antibody antigen complexes, see targets 233 and 234) the structural models of the unbound monomers remain highly valuable for integrative modeling purposes. In the following we discuss in more details a few notable targets.

Target 160 (see Fig. 1A) was unusual to work on, for the reason that a hint was published online. Indeed, as the main strength of HADDOCK is coming from its ability to drive the docking using information, having access to a Twitter-published picture of the complex enabled us to make measurements and derive restraints to guide the docking based on this data. This showcases how broad the spectrum of data can be used in HADDOCK and that one should not be limited in its definition of what kind of information can be provided. Despite none of the interfaces being of acceptable quality, the overall shape of the complex was well modeled. Another aspect worth discussing for this target concerns the accuracy of the nanobody loop conformations, a crucial aspect when performing docking. The predicted nanobody loops were notably poor, with the majority of structures showing a CDR RMSD (calculated with respect to the bound structure [31] after aligning over the framework regions) higher than 5 Å. More specifically, the H3 loop of chain H (nanobody694) was completely mispredicted for all the MODELLER models (CDR RMSD around 10 Å), in which the extended H3 loop conformation was predicted rather than the kinked one. Modern deep learning-based methods generally predict this germline-dependent conformation more accurately [32].

Target 231 (see Fig. 1C) is an interesting case for us. In this case the 3_10_-helical conformation of the peptide was not modeled correctly, but HADDOCK was still capable of generating models sharing a good fraction of native contacts with the published structure [33]. Another lesson to learn here concerns the hypervariable region of the antibody: the CDR RMSDs (1.24 Å and 1.02 Å) between our acceptable models and the antibody in the bound form (PDB ID 8RM0) are higher than the one calculated (0.49 Å) between the bound and unbound (PDB ID 7BG1) structure, namely our input antibody. The interface refinement in this case decreased the accuracy of the antibody model, which can happen especially if the starting conformation is already very close to the bound form.

For targets 233 and 234, the prediction of the peptide in the MHC groove was surprisingly not part of the modeling task, but knowledge about its presence would have been useful to filter out the models in which one or more CDR loops were overlapping with the typical position of peptide in a peptide-MHC complex.

For Target 234 we were the only group capable of identifying a good model for this complex in the scoring challenge. Interestingly, we generated two near-acceptable poses in the prediction challenge, namely the second-best model in terms of Fnat (*F*_*nat*_ = 0.36) and the second-best model in terms of Ligand-RMSD (L-RMSD = 10.6 Å) across all the predictors. The former is important in the context of antibody design, where the correct identification of the interface contacts may play a more important role than the correct modeling of the complex (as given by interface and ligand RMSD). Regarding the scoring, the selected complex was one of the four acceptable-or better models among the shuffled CAPRI scoring ensemble consisting of 1677 structures. We discovered that the model was generated by us and placed in position 85 of the decoy set we provided for the scoring challenge. This indicates that our scoring during the prediction stage could be improved.

## 4 Conclusions

CAPRI rounds 47-55 featured very challenging targets, the majority of which involved antibody/nanobody antigen interactions. Although a significant improvement in the modeling of such interactions has been observed in the last years [35–37], the accurate prediction of an antibody-antigen interface when no experimental information is available is still a challenging task.

While HADDOCK faced difficulties in these CAPRI rounds, it successfully identified accurate models for a couple of difficult targets. The transition from mainly classical, physics-based approaches to deep-learning-aided integrative modeling strategies, where AlphaFold2 and other machine-learning tools provide input structures (and sometimes even interface restraints) represents a paradigm shift for the docking community and should certainly help reaching higher success rates in the future CAPRI challenges. We have seen new methods recently appearing, such as ColabDock [38], that also make use of contact restraints to guide the modeling process, that could be the next generation of deep-learning docking strategies, but yet still limited to protein-protein interactions.

Development and testing of the new version of HADDOCK (v3.0) are currently ongoing and we expect that the addition of new modules, both for the docking and refinement stages as well as for scoring and analysis, will enhance HADDOCK’s ability to generate and properly rank higher quality models.

## 5 Data Availability Statement

All the data reported in this paper can be found at the CAPRI assessment webpage: www.capri-docking.org/assessment

## 6 Acknowledgements

This project has received funding from the European Union Horizon 2020, projects BioExcel (823830 and 101093290) and EGI-ACE (101017567), and from the Netherlands e-Science Center (027.020.G13).

