## Supplementary information for "Integrative modeling in the age of machine learning: a summary of HADDOCK strategies in CAPRI rounds 47-55" (from another version of this preprint: version 1, 2024-09-19)

### 1 HADDOCK3 scoring example

Here we report an example prototype of the HADDOCK 3.0 scoring workflow executed for the majority of the targets. First, the topology of the models is built thanks to the `topoaa` module. Second, a short energy minimisation is carried out with the OPLS force field [1] (`emscoring` module). Then, Fraction of Common Contacts clustering [2] (`clustfcc`) is carried out with default parameters, the most important one being the `clust_cutoff` (0.6 by default). If several good scoring models are labelled as "unclustered", (i.e. they belong to a cluster with population lower than 4), the parameter `min_population` can be decreased. `caprieval` follows, in which the remaining models are compared against a reference or, by default, against the best-scoring one. The final step, `seletopclusts`, selects the top 4 models of the first 10 clusters for visualization and submission.

```
# HADDOCK3 scoring example
run_dir = "scoring_t234
molecules="scoring_t234.pdb"

[topoaa]

[emscoring]

[clustfcc]

[caprieval]

[seletopclusts]
```
